# A Bayesian Multivariate Mixture Model for Spatial Transcriptomics Data

**DOI:** 10.1101/2021.06.23.449615

**Authors:** Carter Allen, Yuzhou Chang, Brian Neelon, Won Chang, Hang J. Kim, Zihai Li, Qin Ma, Dongjun Chung

## Abstract

High throughput spatial transcriptomics (HST) is a rapidly emerging class of experimental technologies that allow for profiling gene expression in tissue samples at or near single-cell resolution while retaining the spatial location of each sequencing unit within the tissue sample. Through analyzing HST data, we seek to identify sub-populations within a tissue sample that reflect distinct cell types or states. Existing methods either ignore the spatial heterogeneity in gene expression profiles, fail to account for important statistical features such as skewness, or are heuristic network-based clustering methods that lack the inferential benefits of statistical modeling. To address this gap, we develop SPRUCE: a Bayesian spatial multivariate finite mixture model based on multivariate skew-normal distributions, which is capable of identifying distinct cellular sub-populations in HST data. We further implement a novel combination of Pólya–Gamma data augmentation and spatial random effects to infer spatially correlated mixture component membership probabilities without relying on approximate inference techniques. Via a simulation study, we demonstrate the detrimental inferential effects of ignoring skewness or spatial correlation in HST data. Using publicly available human brain HST data, SPRUCE outperforms existing methods in recovering expertly annotated brain layers. Finally, our application of SPRUCE to human breast cancer HST data indicates that SPRUCE can distinguish distinct cell populations within the tumor microenvironment.

## 1 Introduction

High throughput spatial transcriptomics (HST) is a developing class of experimental technologies that has proven invaluable in studying a wide range of biological processes in both diseased (van den Brink et al., 2020; Chen et al., 2020) and healthy (Baccin et al., 2020; Mantri et al., 2020) tissues. The advantage of HST over existing sequencing tools like single-cell RNA-sequencing (scRNA-seq) is that HST preserves the spatial location of cells within a tissue sample, while scRNA-seq decouples gene expression information from cell locations during the sequencing process (Burgess, 2019). However, since spatial proximity has been shown to be a principal source of heterogeneity in important biological settings such as the tumor microenvironment (Janiszewska, 2020; Moncada et al., 2018), it is critical to properly weigh both the spatial location of cells and their gene expression profiles when analyzing HST data.

Since the advent of HST technologies, a small number of computational and statistical methods have been proposed to analyze spatially resolved gene expression data to infer biologically distinct sub-populations of cells within a tissue sample – a critical and foundational step in the analysis of HST data. Dries et al. (2019) introduced Giotto, a nearest neighbors network-based clustering tool that offers the ability to cluster cells based on gene expression information only using the Louvain algorithm (Blondel et al., 2008), then spatially refine cell cluster assignments using a hidden Markov random field model. Similarly, in a recent version of the popular scRNA-seq analysis package Seurat, Hao et al. (2020) included the ability to incorporate spatial information into the cell clustering using a spatially-weighted similarity matrix. In a related work, Pham et al. (2020) proposed stLearn, which clusters cells by applying the Louvain or K-means algorithm to a spatially perturbed dimension reduction of the gene expression space, then infers spatial sub-clusters using the DBSCAN algorithm (Ester et al., 1996). While these methods offer the ability to introduce spatial information into standard cell clustering routines, they each adopt network-based approaches that depend heavily on tuning parameters like the number of neighbors and cell clustering resolution, and thus lack the inferential benefits of statistical modeling, such as uncertainty quantification and optimization of parameters using model fit criteria.

Zhao et al. (2021) improved on these works by developing BayesSpace, a Bayesian multivariate-*t* mixture model that induces spatial correlation in mixture component assignment by placing a Potts model prior on mixture component weights. However, BayesSpace is limited in that (i) it models principal components of gene expression features instead of directly modeling gene expression, thus reducing the interpretability of results and obfuscating the need for a multivariate approach since principal components are, by definition, orthogonal (Abdi and Williams, 2010); (ii) BayesSpace assumes symmetric multivariate outcome distributions, which makes its direct application to gene expression features difficult to justify, due to the inherent skewness of gene expression across a tissue sample as shown in Section 2; (iii) BayesSpace uses a global spatial smoothing parameter that must be chosen *a priori* to induce spatial correlation, thus ignoring important local heterogeneities in spatial patterns across a tissue sample; and (iv) while BayesSpace enforces spatially smooth mixture component assignments, gene expression features are assumed to have no spatial dependence after conditioning on mixture component labels – a simplifying assumption that precludes modeling spatial heterogeneity within mixture components.

To address these gaps, we developed SPRUCE (**SP**atial **R**andom effects-based cl**U**stering of single **CE**ll data) for robust identification of cell type sub-populations using HST data. Our proposed model extends the current methodology in a number of ways. First, SPRUCE models gene expression features directly using a multivariate approach. By doing so, we allow for more natural interpretation of mixture components as sub-groups of cells with distinct transcriptional regulatory factors (Wan et al., 2019), and thus distinct gene expression profiles. Next, while existing approaches consider spatial information only in the cluster allocation portion of the mixture model, SPRUCE directly accounts for spatial dependence in both gene expression outcomes and cell-type membership probabilities using spatially correlated random effects in each model component. This model design allows for local heterogeneities in gene expression that can be explained by spatial information, and offers the ability to infer spatially smooth mixture components across a tissue sample. We also accommodate skewed gene expression distributions – a ubiquitous feature of HST data that results from normalization of overdispersed RNA read counts. Finally, SPRUCE relies on a robust and efficient Gibbs sampling algorithm with built-in protection against label switching. SPRUCE is implemented using a novel application of Pólya–Gamma data augmentation to allow for Gibbs sampling of all model parameters, thereby improving upon the reliability of existing methods.

## 2 Data

While a variety of experimental methods exist for measuring spatially resolved RNA abundance in a tissue sample, we focus on high throughput sequencing-based technologies such as the popular 10X Genomics Visium, which allow for measurement of the entire transcriptome instead of a smaller subset of pre-specified genes. Current sequencing-based HST technologies divide the tissue sample into a contiguous array of “spots”, each roughly 55 *μ*m in diameter and containing a small number (often < 5) of spatially close cells (Maniatis et al., 2021). *In situ* barcoding of spots is then used to correlate spatial centroids with the expression levels of thousands of RNAs in each spot (Maniatis et al., 2021). Raw sequencing-based HST data take the form of (i) a spot-by-gene expression matrix, where the number of spots is between 1,000 to 5,000 and the number of genes can exceed 30,000 in most samples (Maniatis et al., 2021); and (ii) a 2-dimensional coordinate matrix locating the centroid of each sequencing spot within the tissue sample. However, it has been shown that there exists vast statistical redundancy in the genes sequenced due to either highly correlated or lowly expressed genes (Edsgärd et al., 2018), and thus we first select a small subset of spatially variable genes (SVGs) using either pre-existing feature selection methods (Hafemeister and Satija, 2019; Edsgärd et al., 2018; Hao et al., 2020) or by focusing on known marker genes.

To illustrate the important characteristics of HST data, we plot in Figure 1 the spatial expression patterns and within-layer densities of three SVGs in a human brain tissue sample (Maynard et al., 2021), in which 33,538 genes were sequenced across 3,085 cell spots using the 10X Visium platform. In Section 5.1, we explore this particular data set in more detail using the expert annotations of brain layers provided by Maynard et al. (2021) as ground truth to benchmark our proposed statistical model relative to existing tools. To quantify the spatial autocorrelation of gene expression throughout the human brain tissue sample, we computed Moran’s I statistic (Gittleman and Kot, 1990; Paradis and Schliep, 2019) and associated p-value for three SVGs identified using standard approaches (Edsgärd et al., 2018), namely PCP4, MBP, and MTCO1. Further, we quantified skewness of gene expression within each expert annotated brain layer using sample skewness (Joanes and Gill, 1998; Meyer et al., 2021).

**Figure 1:**
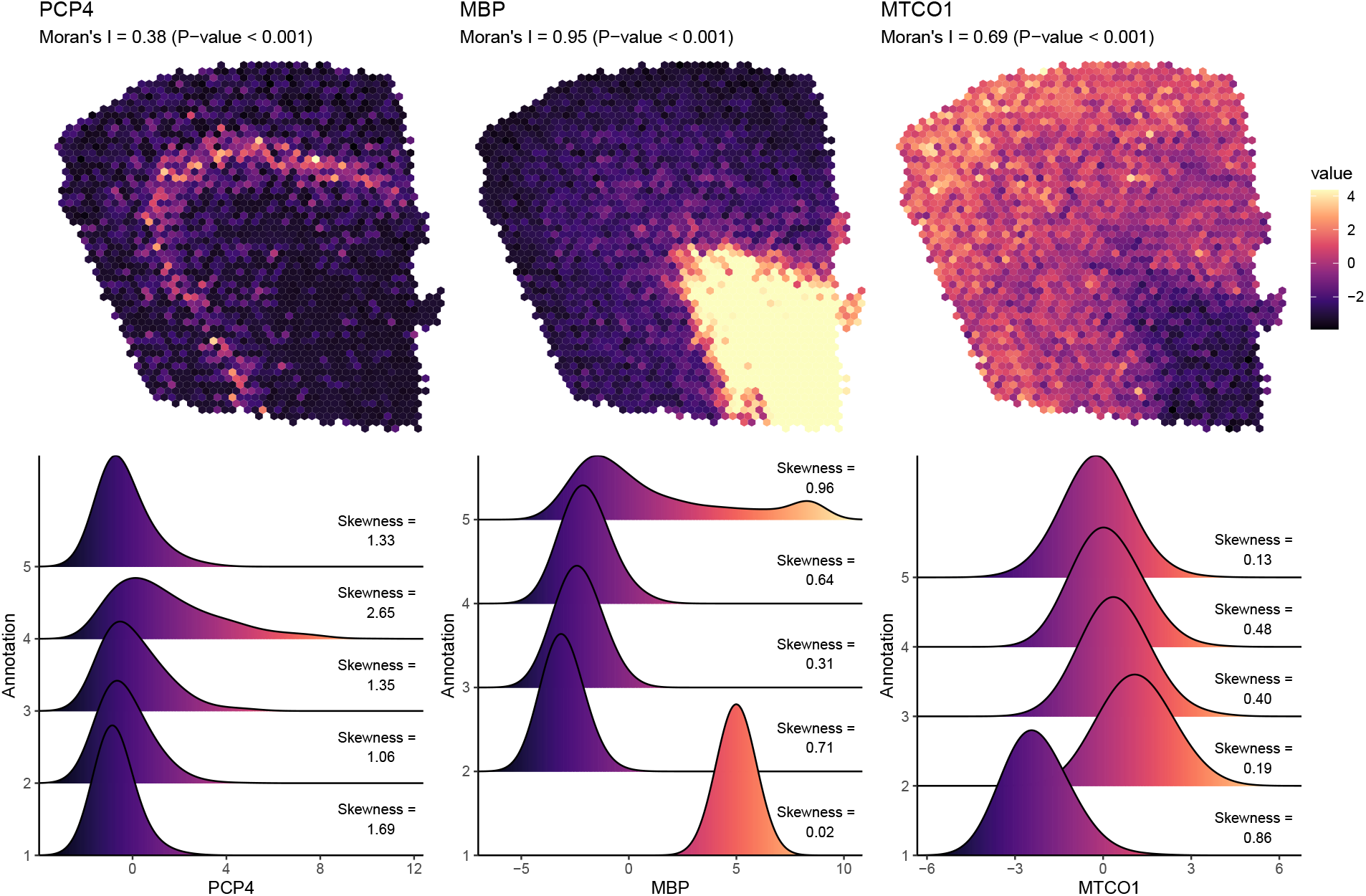
Human brain slice sequenced with the 10X Genomics Visium platform. (Top row) The spatial expression patterns of three SVGs: PCP4, MBP, MTCO1 are shown across the tissue sample, where brighter colored spots correspond to higher expression. Moran’s I statistics and associated p-values suggest significant spatial autocorrelation present in gene expression. (Bottom row) Gene expression densities are shown for each gene within each expert annotated tissue layer along with sample skewness statistics. Empirical densities and skewness statistics imply the need for accommodation of non-symmetric gene expression features.

As shown in Figure 1, the expression of certain genes across a tissue sample can exhibit high spatial variability, hence the need for robust statistical models that account for spatial correlation in gene expression. In addition to spatial correlation, Figure 1 shows residual skewness of gene expression features after accounting for brain tissue region using expert annotations. In fact, skewness occurs in most all normalized gene expression features due to the nature of converting overdispersed count data to normalized data. Thus, any robust statistical model for HST data analysis should allow for non-symmetric gene expression distributions.

While the human brain is a natural choice for benchmarking HST data analysis methods due to its well-studied spatial structure, it is of great scientific need to generate similar insights in settings like the breast cancer tumor microenvironment – a disease area that has yet to be studied using existing HST data analysis tools. To address this gap, in Section 5.2 we analyzed the human invasive breast cancer tumor sample made publicly available by 10X Genomics (10x Genomics, 2020), which consists of 36,601 genes measured across 3,798 spots generated by the 10X Visium platform. Figure 4A shows the spatial expression patterns and densities of a selection of SVGs across the breast tissue sample. Here, we see that the strength of spatial autocorrelation differs between the human brain tissue sample and the breast cancer tumor sample, hence the need for allowing spatial information to enter flexibly into our statistical model for HST data.

## 3 Model

In Section 3, we present SPRUCE, a Bayesian spatial mixture model capable of addressing the important challenges presented by HST data described in Section 2. First, in Section 3.1, we develop a general multivariate mixture model framework that is capable of clustering cells while accounting for spatial correlation, gene-gene correlation, and skewness of gene expression features. Then, in Section 3.2 we improve upon previous approaches for analyzing HST data by implementing a novel cluster-membership model that combines Pólya–Gamma data augmentation with spatially-correlated random effects to induce spatial dependence among neighboring cells and allow for robust interpretation of mixture components. Section 3 concludes with discussion of prior distributions, accommodation of heavy-tailed gene expression features, model selection, and Markov chain Monte Carlo (MCMC) simulation details.

### 3.1 General Mixture Model

Our proposed model is relevant for sequencing-based HST platforms such as 10X Visium, which, as shown in Figure 1, divide the tissue sample into a regular lattice of cell spots. Each spot is associated with a high-dimensional gene expression profile that can be used to infer cell type (e.g., T cells, B cells, natural killer cells, *et cetera* in the context of immune cells) (Asp et al., 2020). To identify these biologically relevant sub-populations within the tissue sample, we adopt a finite mixture model that accounts for important features of the data such as spatial dependence among cell spots, dependence across correlated genes, and non-normality of gene expression profiles. Our approach extends existing spatial finite mixture models in the statistical literature (Neelon et al., 2014) to this challenging setting.

Let **y***_i_* = (*y*_*i*1_, …, *y_ig_*)*^T^* be the length *g* vector of gene expression features for spot *i*(*i* = 1, …, *n*). As discussed in Section 5, the standard pre-processing steps for HST data include identification and normalization of the *g* top SVGs before modeling (Edsgärd et al., 2018). To identify biologically relevant cell sub-populations within a tissue sample using these *g* pre-selected spatially variable gene expression features, we propose a finite mixture model of the form

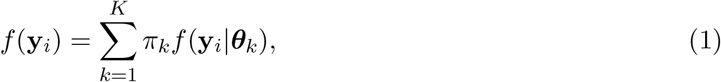

where ***θ**_k_* is the set of parameters specific to component *k*(*k* = 1, …, *K*) and π*_k_* is a mixing weight that measures the probability of a given spot belonging to cell sub-population *k*. For now, we assume **π** = (π_1_, …, π*_k_*) is common to all cell spots, though in Section 3.2 we develop a model for cell spot-specific mixing weight parameters. The number of cell sub-populations *K* may be specified based on biological knowledge, or may be identified entirely from the data, as described in Section 3.4.2.

To facilitate Bayesian inference, we introduce latent cluster indicator variables *z*_1_, …, *z_n_*, where *z_i_* ∈ {1, …, *K*} indicates the mixture component assignment for cell spot *i*. Given *z_i_* = *k*, we assume that the gene expression features for spot *i* follow a *g*—dimensional multivariate skew normal (MSN) distribution (Azzalini and Valle, 1996)

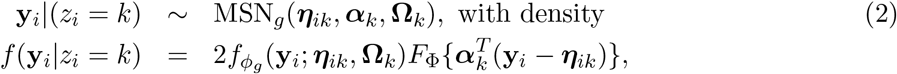

where, given *z_i_* = *k*, ***η**_ik_* is the length *g* mean vector for spot *i*, ***α**_k_* is a length *g* vector of feature-specific skewness parameters for mixture component *k*, **Ω**_*k*_ is a *g* × *g* scale matrix that captures association among the gene expression features in mixture component *k*, *f_ϕ_g__*(**y***_i_*; ***η**_ik_*, **Ω**_*k*_) is the density function of a *g*-dimensional normal distribution with mean ***η**_ik_* and variance-covariance matrix **Ω**_*k*_ evaluated at **y**_*i*_, and *F*_Φ_ is the CDF of a scalar standard normal random variable. When ***α**_k_* = **0**_*g*×1_, the distribution of **y**_*i*_ is multivariate normal (MVN) with mean ***η**_ik_* and variance-covariance matrix **Ω**_*k*_. Positive elements of ***α**_k_* imply positive skewness relative to the MVN distribution, while negative values imply negative skewness. While model (2) allows for mixture component and feature-specific departures from normality in terms of skewness, in Section 3.3 we further extend model (2) to accommodate heavy tailed gene expression densities using the multivariate skew-*t* distribution.

We may represent the MSN distribution using a convenient conditional representation in terms of the MVN distribution and a spot-level standard normal random variable truncated below by zero *t_i_* ~ N_[0,∞)_(0,1) (Frühwirth-Schnatter and Pyne, 2010). To implement this conditional MSN representation and incorporate spatial variability across the tissue sample into the gene expression model, we let

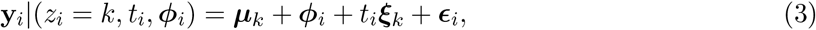

where ***μ**_k_* is the length *g* gene expression mean vector for mixture component *k*, ***ϕ**_i_* is a length *g* spatial effect that allows for spatially-correlated departure from ***μ**_k_* in spot *i*, ***ξ_k_*** controls the mixture component-specific skewness of each gene expression feature in the conditional MSN representation, and ***ϵ_i_*** ~ N_*g*_ (**0**, **Σ***_k_*). In Web Appendix B, we describe how the original MSN parameters ***η**_ik_*, ***α**_k_*, and **Ω**_*k*_ can be obtained through back-transformations as functions of the parameters in equation (3).

To accommodate spatial dependence among cell spots in the tissue sample, we adopt a multi-variate intrinsic conditionally autoregressive (CAR) prior (Besag, 1974) for ***ϕ**_i_*:

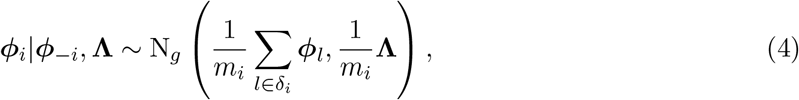

where ***ϕ**_−i_* denotes the spatial random effects for all spots except spot *i*, **Λ** is a *g* × *g* variance-covariance matrix for the elements of ***ϕ**_i_*, *m_i_* is the number of neighbors of spot *i*, and *δ_i_* is the set of all neighboring spots to cell spot *i*. To aid in separability between **Λ** and **Σ**_*k*_, we assume the variance-covariance of the spatial random effects **Λ** is shared across mixture components, while **Σ**_*k*_, the conditional variance-covariance of **y**_*i*_, is mixture component-specific. As stated by Brook’s lemma (Banerjee et al., 2014), model (4) leads to a uniquely defined yet improper joint distribution of (***ϕ***_1_, …, ***ϕ**_n_*):

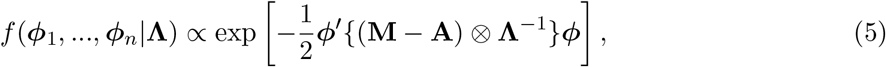

which is due to the fact that the *n* × *n* matrix (**M** – **A**) is singular, where **M** = diag(*m*_1_, …, *m_n_*), **A** is the *n* × *n* adjacency matrix for all cell spots within the tissue sample, with *A_ij_* = 1 if cell spot *i* borders cell spot *j* and *A_ij_* = 0 otherwise, and ***ϕ*** is a length *gn* vector formed by concatenating ***ϕ***_1_, …, ***ϕ**_n_*. However, as described in Banerjee et al. (2014), we ensure a proper posterior distribution for each ***ϕ**_i_* by enforcing a sum-to-zero constraint on the elements of each ***ϕ**_i_* for *i* = 1, …, *n*. In Section 3.4.1, we complete the fully Bayesian model specification by assigning conjugate priors to all remaining model parameters, thus leading to closed-form full conditional distributions for all model parameters and allowing for an efficient Gibbs sampling algorithm detailed in Web Appendix B.

The SPRUCE model presented in this section for the analysis of sequencing-based HST data has several desirable features and provides distinct advantages over existing methods. First, through the use of spatially correlated random effects, the spot-level SPRUCE model explicitly accounts for spatial variability of gene expression throughout a tissue sample that is not explained by cell type (i.e., ***μ**_k_*). Next, SPRUCE directly accommodates skewness in each mixture component density - a common feature in gene expression data that is ignored by existing methods for clustering single cell data which assume symmetric distributions of gene expression features (Zhao et al., 2021). Finally, SPRUCE provides the inferential benefits of a fully Bayesian approach, such as the ability to make posterior probability statements about all model parameters and the ability to choose *K*, the number of sub-populations, in a principled and model-based manner as described in Section 3.4.2.

### 3.2 Spatial Pólya–Gamma Multinomial Logit Regression Component Membership Models

Thusfar, we have assumed that spatial dependence enters only into the model for gene expression distributions, where each spot is allowed to vary with respect to its mixture component-specific mean through the use of spatially correlated multivariate random effects. However, in many cases we may wish to allow the probability **π** of belonging to each mixture component to vary spatially as well. In doing so, we may ensure that the cellular sub-populations identified by the model are informed by the spatial variability across tissue samples. First, we extend model (1) by letting

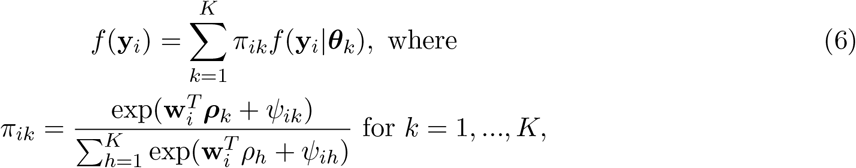

where π*_ik_* = P(*z_i_* = *k*), **w***_i_* is a length *p* vector of covariates relevant to cluster membership, ***ρ**_k_* is an associated length *p* vector of fixed-effects, and *ψ_ik_* is a spatial random effect allowing spatially-correlated variation with respect to 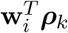. For identifiability purposes, we choose mixture component 1 as the reference category and set ***ρ***_1_ = **0**_*p*×1_ and *ψ*_*i*1_ = 0 for all *i* = 1, …, *n*. To introduce spatial association into the component membership model, we assume univariate intrinsic CAR priors for *ψ_ik_*:

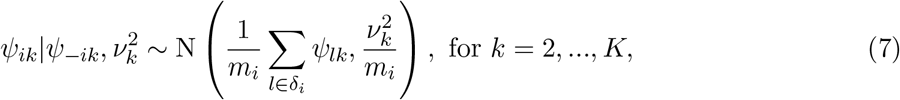

where 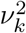 is a mixture component-specific variance for *ψ_ik_*.

We ensure closed-form full conditional distributions of the multinomial logit regression parameters by adopting a Pólya-Gamma data-augmentation approach as introduced by Polson et al. (2013). In the context of Bayesian logistic regression, Polson et al. demonstrate that the inverse-logit function can be expressed as a scale-normal mixture of Pólya-Gamma densities, and the likelihood of the logistic model can in turn be written as a scale-mixture of normal densities, allowing for closed-form conditional distributions of all model parameters. While previous models (Allen et al., 2020) have applied these results from Polson et al. for use in multinomial logit mixture weight regression models, the Pólya–Gamma data augmentation approach has yet to be used in conjunction with CAR priors in the context of modeling mixing weights in spatial finite mixture models. In Proposition 1 below, we show that Pólya–Gamma data augmentation allows for closed-form full conditional distributions of *ψ_ik_* in this novel setting.

#### Proposition 1

*Let π_ik_ follow the multinomial logit model defined in equation (6), and let ψ_ik_ have a univariate intrinsic CAR prior as defined in equation (7). Under Pólya–Gamma data augmentation, the full conditional distribution of ψ_ik_ is N(m_ik_, V_ik_), where*

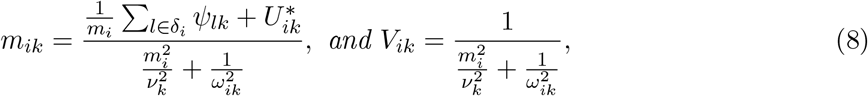

*where* 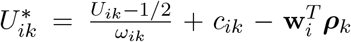, *U_i,k_ is an indicator equal to* 1 *if z_i_* = *k and* 0 *otherwise*, 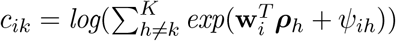, *and ω_ik_ ~ PG*(1, 0). *The proof is provided in Web Appendix A*.

### 3.3 Extensions to Multivariate Skew-*t* Distributions

In the case of outliers or heavy-tails in the distributions of gene expression features, we extend model (2) to the multivariate skew-*t* (MST) distribution (Gupta, 2003):

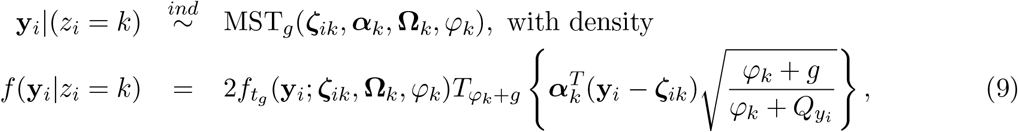

where *f_t_g__* (**y**_*i*_; ***ζ**_ik_*, **Ω**_*k*_, *φ_k_*) denotes the CDF of a *g*-dimensional *t* distribution with location ***ζ**_ik_*, covariance **Ω**_*k*_, and fixed degrees of freedom *φ_k_* that may vary across mixture components; *T_φk +g_* denotes the distribution function of the scalar standard *t* distribution with *φ_k_ + g* degrees of freedom; and 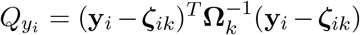. Similarly to the MSN distribution, we may adopt a convenient conditional representation for the MST distribution in terms of standard densities to allow for straightforward Gibbs sampling in the MST setting (Frühwirth-Schnatter and Pyne, 2010). For inference with heavy-tailed gene expression features, we extend model (3) as

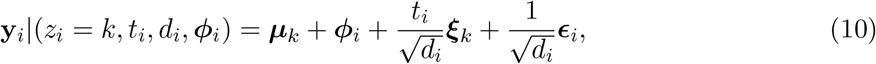

where *d_i_*|(*z_i_* = *k*) ~ Gamma(*κ_k_*/2, *κ_k_*/2) is a spot-specific scale term, and *κ_k_* is a pre-specified degrees of freedom for each mixture component *k* = 1, …, *K*. Lower values of *κ_k_* allow for heavier tails relative to MVN in cluster *k*.

### 3.4 Bayesian Inference

#### 3.4.1 Priors

We complete a fully Bayesian specification of the SPRUCE model by assigning prior distributions to all remaining model parameters. For *k* = 1, …, *K*, we assign cluster specific priors 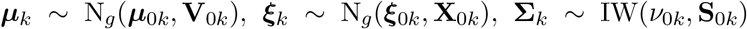. By default, we opt for weakly-informative priors (Gelman et al., 2013) by choosing 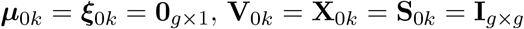, and *ν*_0*k*_ = *g* + 2, which gives *E*(**Σ**_*k*_) = **I***_g×g_*. We further assume **Λ** ~ IW(λ_0_, **D**_0_) for *k* = 1, …, *K*. Weakly-informative priors result from setting λ_0_ = λ_0*k*_ = *g* + 2, and **D**_0_ = **D**_0*k*_ = **I***_g×g_*. Finally, for *k* = 2, …, *K*, we assume ***ρ**_k_* ~ N*_p_*(***ρ***_0*k*_, **R**_0*k*_) and 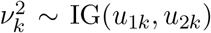, where we obtain weakly-informative priors by choosing ***ρ***_0*k*_ = **0**_*p*×1_, **R**_0*k*_ = **I***_p×p_*, and *u*_1*k*_ = *u*_2*k*_ = 0.001. A detailed description of the resultant Gibbs sampling algorithm is provided in Web Appendix B.

#### 3.4.2 Model Selection

The choice of *K*, i.e., the number of mixture components used in the SPRUCE model, is a critical step in the analysis of HST data. In some situations, it may be appropriate to specify *K* based on strong biological knowledge of the cell types that will be present in a tissue sample, or the desire to investigate a known number of “cell states” within a more homogeneous tissue sample. In other cases, however, such prior information might be unavailable and the choice of *K* should be made entirely based on the data. To identify the optimal value of *K* in terms of model fit, we make use of the widely applicable information criterion (WAIC) (Watanabe, 2010) defined as

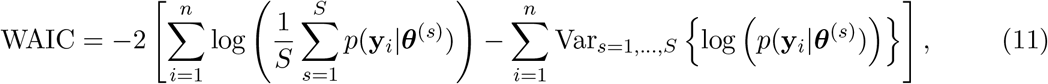

where *s* = 1, …, *S* indexes the post-burn-in iterations of the Gibbs sampler detailed in Web Appendix B, and let ***θ***^(*s*)^ represents the current values of all parameters at iteration *s*.

#### 3.4.3 Label Switching

Label switching is a common issue faced by Bayesian mixture models in which the invariance of the likelihood to permutations of **z** = (*z*_1_, …, *z_n_*) results in conflation of cluster-specific parameters across MCMC iterations due to statistically equivalent permutations of **z** (Stephens, 2000; Jasra et al., 2005). Existing approaches for addressing the label switching issue either attempt to re-shuffle posterior samples after MCMC convergence (Papastamoulis, 2016) or impose an arbitrary order restrictions on the component-specific parameters ***θ***^(*s*)^. However, these existing approaches are not ideal since (i) re-shuffling of posterior samples relies on prediction of component label re-mappings, thereby introducing the potential for additional error that may impede the accuracy of component-specific parameter estimates; and (ii) imposing order constraints on ***θ**_k_* can lead to poorly estimated parameters when mixture components are not well-separated.

To overcome these challenges, we adopt the “canonical” remapping approach proposed by Peng and Carvalho (2016) in the context of network community detection using blockmodels. Here, Bayesian inference relies on sampling discrete community indicators, and thus is similarly susceptible to the label switching problem. Peng and Carvalho avoid this issue by restricting the sample space of **z** to a canonical sub-space, and define a canonical projection to remap the sampled **z** at each MCMC iteration to the canonical sub-space. The canonical sub-space 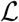 is defined as 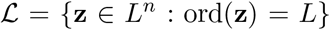, where *L* = (1, …, *K*), and ord(**z**) returns the length *K* vector of the order in which each mixture component *k* = 1, …, *K* appears in the vector **z**. Here, ord(**z**)[1] = *z*_1_ is the first unique mixture component to occur in **z**, ord(**z**)[2] is the second unique mixture component to occur in **z**, *et cetera*. We further define 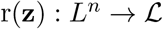 as the canonical projection used to remap **z** to the canonical sub-space at each MCMC iteration as described in Web Appendix B. Finally, we choose as our final estimate of **z** the maximum *a posteriori* (MAP) estimate of of **z** across all post burn-in MCMC samples. This approach is desireable as it does not rely on posterior re-shuffling and does not place restrictions on the within-cluster parameters ***θ***^(*s*)^.

## 4 Simulation Study

To investigate the performance of SPRUCE and validate our proposed Gibbs sampling estimation algorithm, we generated simulated HST data mimicking a publicly available sagital mouse brain data set sequenced with the 10X Visium platform and made available by 10X Genomics (10x Genomics, 2019). To ensure our simulation study is reflective of real HST data sets, we first allocated the *n* = 2, 696 cell spots in the original sagital mouse brain data set into one of *K* = 4 simulated ground truth tissue segments that resemble distinct mouse brain layers (Figure 2A). We then simulated spatially variable multivariate gene expression features of dimension *p* = 16 according to the MSN model with parameters shown in Table 1. Parameters were chosen to result in weakly separated mixture components, as is shown by the Uniform Manifold Approximation and Projection (UMAP) (McInnes et al., 2018) dimension reduction in Figure 2B. Next, we fit three model variants: (i) an MVN mixture model with no spatial random effects; (ii) an MSN mixture model with no spatial random effects; and (iii) an MSN mixture model with spot-level multivariate CAR spatial random intercepts in the gene expression model. This set of models allows us to demonstrate how accounting for skewness and spatial correlation in gene expression outcomes may lead to improved parameter estimates relative to ground truth. Each model was run for 10, 000 MCMC iterations, with the first 1, 000 iterations discarded as burn-in, and priors were chose to be weakly informative as described in Section 3.4.1.

**Figure 2:**
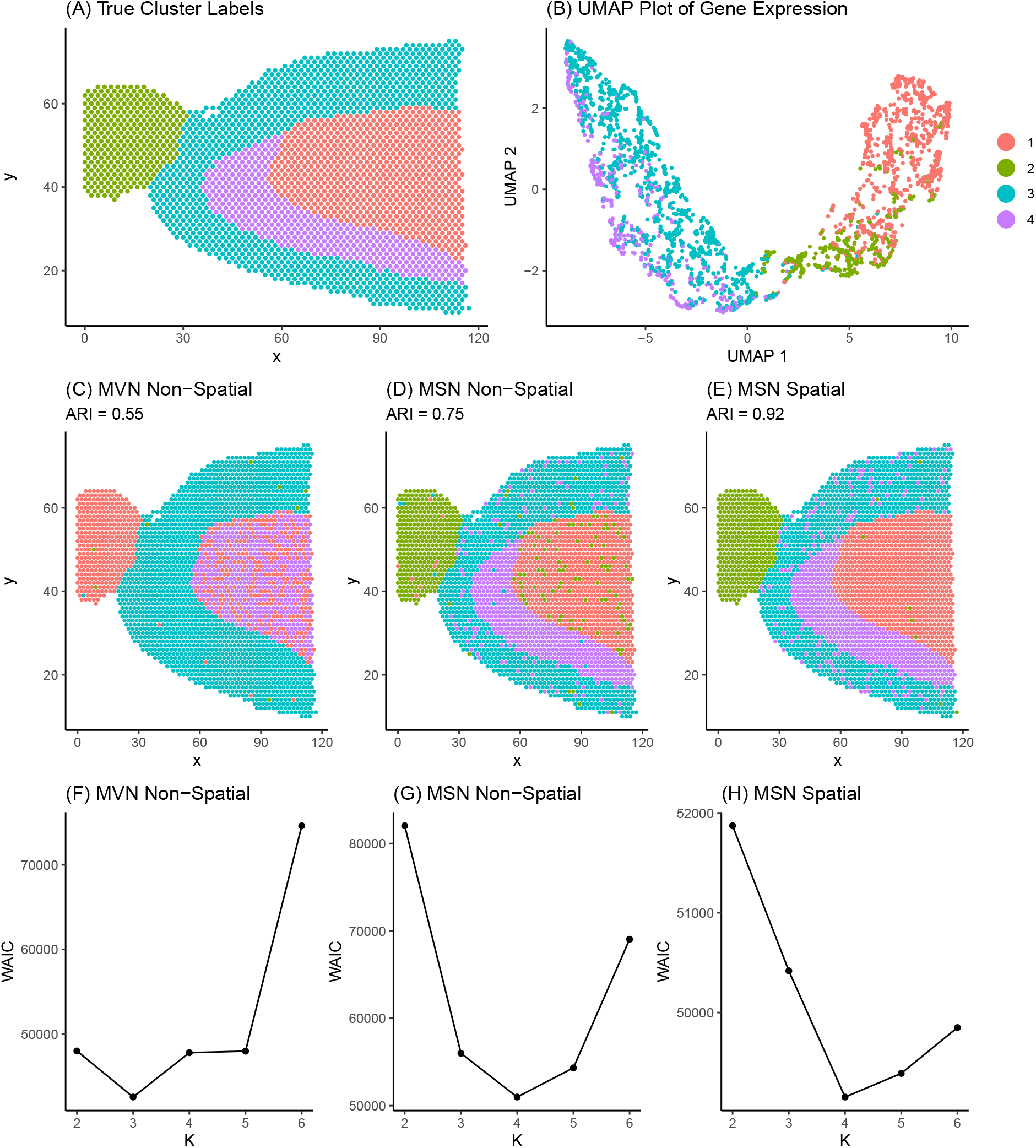
Sagital mouse brain tissue sample manually segmented into four regions. (A) True simulated cluster labels. (B) UMAP dimension reduction of simulated gene expression matrix. Points correspond to tissue spots in the sagital mouse brain. Points are colored according to ground truth cluster labels and are positioned in the 2-dimensional UMAP space according to their similarity in gene expression. (C) - (E) Model estimated cluster labels. (F) - (H) WAIC model selection curves.

**Table 1:** Simulated parameter values and estimates obtained from three model variants: (i) MVN: multivariate normal clustering without spatial random effects; (ii) MVN Spatial: multivariate normal clustering with CAR spatial random effects; and (iii) MSN Spatial: multivariate skew-normal clustering with CAR spatial random effects. Parameter estimates are shown as posterior means with associated 95 % credible intervals.

| Parameter | True | MVN | MSN | MSN Spatial |
| --- | --- | --- | --- | --- |
| $\mu_{11}$ | -2.00 | -2.72 (-2.86, 1.77) | -2.14 (-2.74, 1.62) | -2.01 (-2.86, -1.55) |
| $\mu_{12}$ | -1.00 | -2.34 (-2.45, 1.01) | -0.83 (-1.27, -0.79) | -0.87 (-1.04, -0.63) |
| $\mu_{13}$ | 1.00 | -0.97 (-1.07, 0.64) | 1.32 (0.85, 1.53) | 1.03 (0.82, 1.24) |
| $\mu_{14}$ | 2.00 | 1.79 (1.66, 2.89) | 2.12 (1.95, 2.36) | 2.03 (1.84, 2.22) |
| $\xi_{11}$ | -1.50 | / | -0.95 (-1.59, -0.28) | -1.64 (-1.82, -1.46) |
| $\xi_{12}$ | -0.75 | / | -0.41 (-0.79, 0.09) | -0.78 (-1.14, -0.51) |
| $\xi_{13}$ | 0.75 | / | 0.63 (0.36, 0.89) | 0.75 (0.43, 0.95) |
| $\xi_{14}$ | 1.50 | / | 1.27 (0.80, 1.52) | 1.43 (0.8, 1.74) |
| $\Sigma_{111}$ | 1.50 | 2.01 (1.78, 2.68) | 1.45 (0.92, 2.21) | 1.57 (1.42, 1.74) |
| $\Sigma_{112}$ | 1.00 | 1.37 (1.17, 2.13) | 1.11 (0.93, 1.75) | 1.13 (0.89, 1.25) |
| $\Sigma_{113}$ | 0.75 | 0.60 (0.42, 1.77) | 0.75 (0.50, 1.45) | 0.78 (0.64, 0.95) |
| $\Sigma_{114}$ | 0.50 | 0.32 (-0.24, 1.22) | 0.42 (0.25, 1.11) | 0.49 (0.39, 0.60) |
| $\Sigma_{122}$ | 1.50 | 1.37 (1.17, 2.13) | 1.59 (0.39, 0.90) | 1.61 (1.41, 1.72) |
| $\Sigma_{123}$ | 1.00 | 0.79 (0.63, 1.74) | 1.05 (0.68, 1.39) | 0.88 (0.64, 1.05) |
| $\Sigma_{124}$ | 0.75 | 0.32 (0.11, 1.26) | 0.82 (0.49, 0.99) | 0.71 (0.54, 0.99) |
| $\Sigma_{133}$ | 1.50 | 1.71 (1.54, 2.13) | 1.41 (1.01, 1.65) | 1.52 (1.24, 1.83) |
| $\Sigma_{134}$ | 1.00 | 0.32 (0.11, 1.26) | 1.03 (0.76, 1.48) | 1.11 (0.54, 2.13) |
| $\Sigma_{144}$ | 1.50 | 1.48 (1.15, 1.70) | 1.87 (1.01, 1.91) | 1.44 (1.30, 1.96) |

Posterior parameter estimates and 95% credible intervals (CrI’s) for a selection of mixture component *k* = 1 parameters are shown in Table 1. In Figures 2C-2E, we show the estimated mixture component labels for each of the three model variants. We quantified the ability of each model to recover ground truth simulated tissue region labels using the adjusted Rand index (ARI) (Hubert and Arabie, 1985). Finally, in Figures 2F-2H we plot model fit as measured by WAIC for each of the three model variants fit across a range of *K* = 2, …, 6 to assess the ability of each model variant to recover the true tissue region labels.

In Figures 2C trough 2E, we see that accounting for skewness and spatial correlation among spots allows for more accurate recovery of true mixture component labels in terms of ARI. In Figures 2F trough 2H, we see that the minimum WAIC value occurs at the correct value *K* = 4 for the two MSN models, but occurs at *K* = 3 for the MVN non-spatial model. Finally, Table 1 displays posterior means and 95% credible intervals for a selection of model parameters in mixture component 1 for each model. Table 1 reveals that he MSN spatial model was able to most accurately estimate the true model parameters, while the MVN and MSN non-spatial models suffered from decreased accuracy in parameter estimates.

## 5 Applications

### 5.1 Analysis of 10X Visium Human Brain Data

To assess the performance of SPRUCE relative to expert annotations and existing methods for clustering HST data, we analyzed the human dorsolateral prefrontal cortex brain data recently published by Maynard et al. (2021), which consisted of 33,538 genes sequenced in 3,085 spots across the tissue sample. We compared SPRUCE to four existing methods, namely BayesSpace (Zhao et al., 2021), stLearn (Pham et al., 2020), Seurat (Hao et al., 2020), and Giotto (Dries et al., 2019). Due to the highly-organized spatial structure of human brain tissue samples and the presence of known marker genes that can be used to delineate distinct layers of the brain, these data can serve as an important benchmark for SPRUCE and existing methods. In this application, we treat the expert annotations from Maynard et al. (2021) as ground truth and use ARI to quantify the agreement between these gold standard annotation and those obtained by SPRUCE and existing tools.

We first implemented the standard Seurat pre-processing pipeline for 10X Visium data (Hao et al., 2020), which includes discarding low quality features, normalizing and scaling gene expression, and computing dimension reductions. For the normalization step, we adopted sctransform: a model-based variance stabilization transformation approach proposed by Hafemeister and Satija (2019). For the dimension reduction step, we used principal component analysis to find the first 128 principal components, then implemented the UMAP dimension reduction algorithm on this set of principal components to facilitate visualization. We used the top 16 SVGs as features for SPRUCE, many of which were found to be layer characterizing genes by Maynard et al. (2021). The number of SVGs was chosen to result in a parsimonious subset of genes, whose expression collectively spanned the spatial domain of the tissue sample. We ran the SPRUCE model MCMC estimation for 10, 000 iterations with a burn in of 1,000. The estimated cluster labels from SPRUCE were taken as the MAP estimate across all saved MCMC samples. Finally, we used default parameter settings for each of the four existing tools.

Figure 3 shows the estimated tissue layer labels from SPRUCE and the four existing HST tools relative to expert annotations. SPRUCE achieved the highest ARI of 0.75 relative to manual annotations, followed by BayesSpace (ARI = 0.55) which struggled discerning layers 4 and 5. The explicit use of layer-specific spatially variable features with SPRUCE as opposed to BayesSpace’s use of principal components computed from all genes may explain the improved performance, as principal components can be affected by low-quality/noise genes while. Additionally, BayesSpace’s use of a global smoothing prior across the entire tissue sample represents a stronger assumption than SPRUCE’s random effects-based approach, which allows for more flexible spatial correlation patterns. The three network-based approaches stLearn, Seurat, and Giotto each performed poorly relative to the manually annotated ground truth labels (ARI = 0.33, 0.29, and 0.24, respectively).

**Figure 3:**
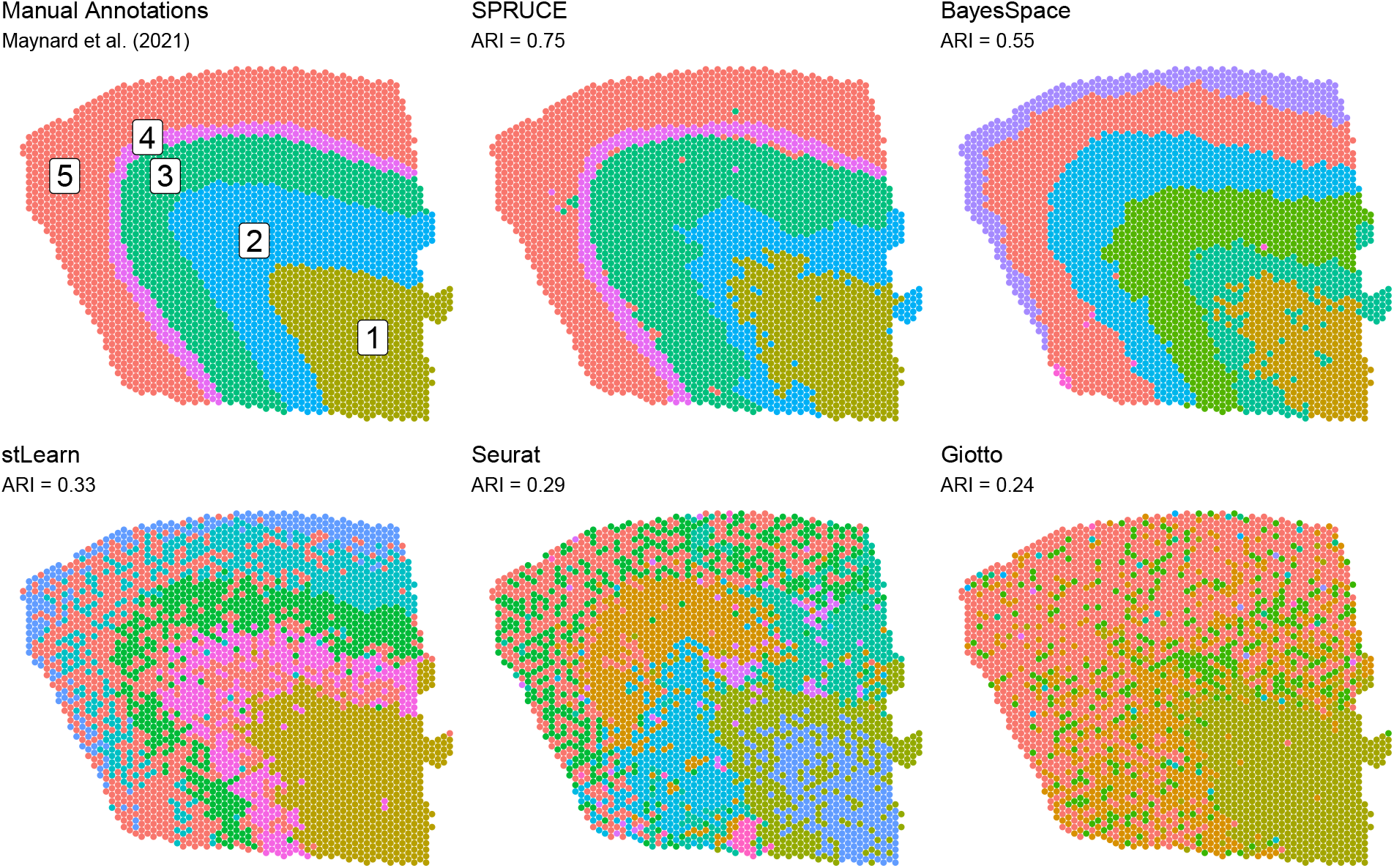
Human brain tissue sample sequenced with the 10X Genomics Visium platform. Expert annotations of brain layers (cell-types) are shown as ground truth labels. ARI measures performance of HST data analysis methods relative to ground truth labels.

### 5.2 Analysis of 10X Visium Breast Cancer Data

To demonstrate the application of our proposed method to the case of unlabeled data, we analyzed a publicly available human Invasive Ductal Carcinoma breast tissue (10x Genomics, 2020) sequenced with the 10X Visium platform. We applied the standard pre-processing pipeline and sctransform normalization approach as in Section 5.1. In Figure 4A, we plot the expression of the top 16 most spatially variable features across the tissue sample. These features display substantial spatial heterogeneity in gene expression, with clear sub-regions existing within the tissue sample. We applied the MSN SPRUCE model with spatially correlated random effects to the normalized breast cancer data, where the 16 top SVGs in Figure 4 were used as features. We identified *K* = 5 as the best fitting model using WAIC and used 10, 000 MCMC iterations with a burn in of 1000.

**Figure 4:**
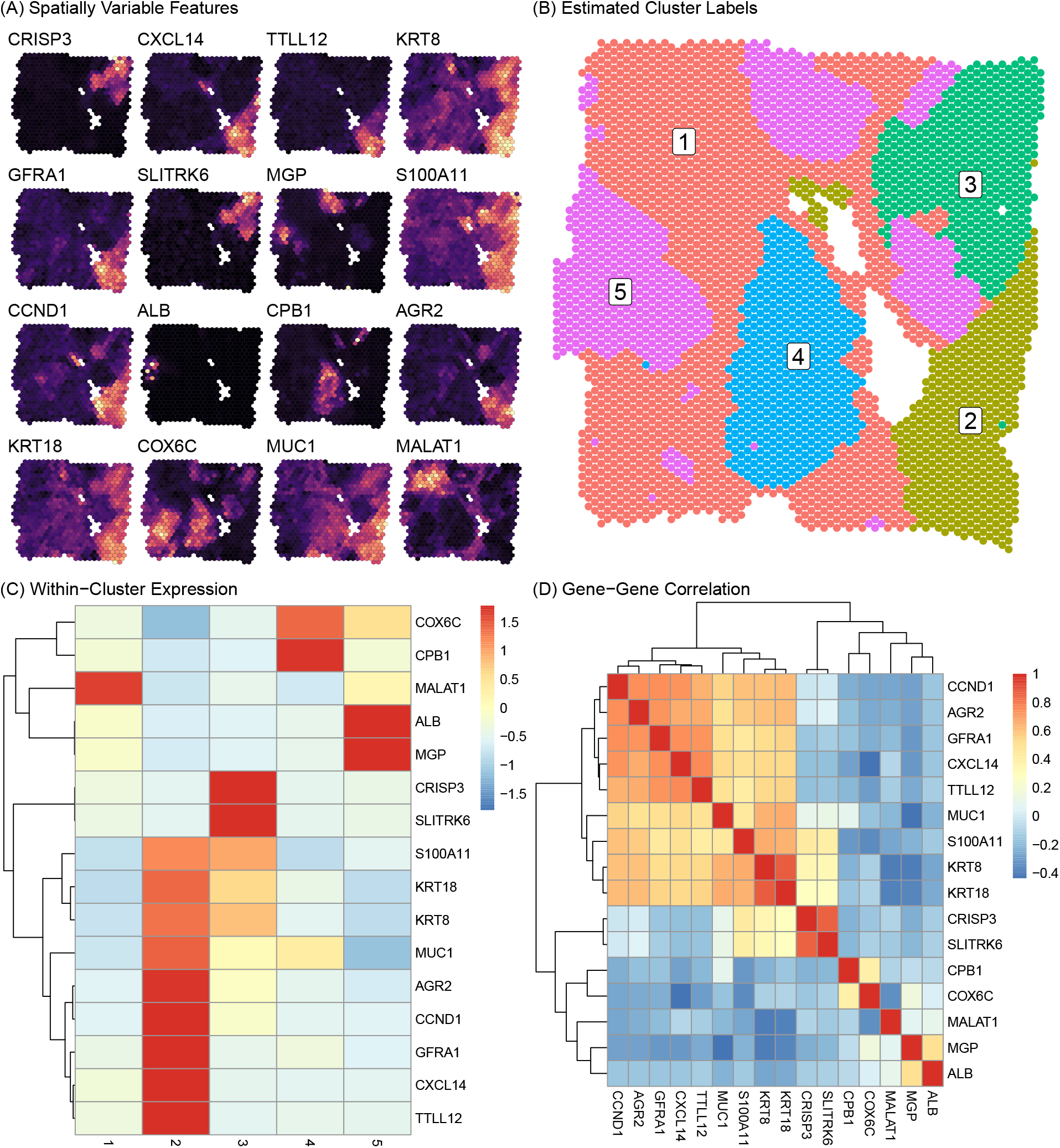
Human Invasive Ductal Carcinoma breast tissue sample sequenced with the 10X Genomics Visium platform. (A) Expression intensity of the top 16 top SVGs is shown across the tissue (brighter color implies higher expression). (B) Inferred cluster labels from SPRUCE. (C) Heatmap of gene mean gene expression profiles within clusters. (D) Heatmap of gene-gene correlations.

Figure 4B shows the MAP estimate of the mixture component labels across the tissue space, which we use to infer distinct sub-regions, i.e., clusters, within the breast tissue sample. To characterize each cluster biologically, we show the posterior mean expression of each gene in each cluster via the heatmap in Figure 4C. This plot shows clearly distinct expression patterns between clusters. Cluster 1 spanned a large portion of the tissue sample and was characterized by medium to low expression of all markers except MALAT1. Cluster 2 was more localized in the bottom right region of the tissue sample and was marked by very high expression of 9 of the 16 genes. This set of 9 genes, as shown in the gene-gene correlation heatmap in Figure 4D, demonstrated highly correlated expression, suggesting a possible pathway function of these genes. Cluster 3 featured high expression of CRISP3 and SLITRK6, but low to moderate expression of all other genes. Similarly clusters 4 and 5 were characterized by high expression of a single pair of genes, namely COX6C and CPB1 in cluster 4, and ALB and MGP in cluster 5.

These results generated by the SPRUCE model may be suggestive of important biological functions related to breast cancer. For instance, expression of MALAT1 has been associated with suppression of breast cancer metastasis (Kim et al., 2018), suggesting cluster 1 may be a region of relatively low tumor expansion within the tissue sample. Meanwhile, cluster 2 expresses tumor-associated antigens (TAAs), i.e., substances produced by tumor cells, such as GFRA1 (Bosco et al., 2018) suggesting cluster 2 as a highly tumor invasive region of the tissue sample. Relatedly, cluster 2 expresses high levels of AGR2, which has been associated with poor breast cancer survival (Ann et al., 2018). Taken together, these results point to an interesting interaction taking place in this breast tissue sample between tumor resistant cells in cluster 1 and cancerous cells in cluster 2. Such findings are illustrative of how SPRUCE may elucidate promising targets for future study across a wide range of disease domains.

## 6 Discussion

We have developed SPRUCE: a fully Bayesian modeling framework for comprehensive analysis of HST, which accounts for important features such as skewness and spatial correlation across the tissue sample. Our model improves upon existing approaches by allowing for a wide range of spatial gene expression patterns via the use of spatially correlated random effects instead of assuming a global smoothing pattern over the tissue sample. We showed how Pólya–Gamma data augmentation can be used to allow for Gibbs sampling of random intercepts modeled with CAR priors in the context of finite mixture model component mixture probabilities. We also established a robust Gibbs sampling algorithm that protects against label switching by remapping mixture component labels to a canonical sub-space.

Through a simulation study based on publicly available 10X Genomics Visium data, we showed how ignoring gene expression features like skewness and spatial correlation can result in poor recovery of true mixture component labels, and bias mixture component-specific parameter estimates. Conversely, when tissue spots are not clearly separated in standard dimension reductions of gene expression features like UMAP, spatial information can be used to help separate distinct sub-populations within the tissue sample. We also showed how model fit criteria such as WAIC may be used to identify the best fitting number of mixture components, which improves upon many existing clustering tools.

We applied SPRUCE to two publicly available 10X Genomics Visium data sets. The first application was concerned with assessing the ability of SPRUCE to recover expert annotations of human brain layers. We found that SPRUCE was best able to discern human brain layers compared to existing methods. Notably, the Bayesian mixture model-based methods (SPRUCE and BayesSpace) performed considerably better than the network-based methods (stLearn, Seurat, and Giotto). We attribute the improved performance of SPRUCE over BayesSpace to the fact that (i) SPRUCE allows for non-symmetry in gene expression features, (ii) SPRUCE models the most spatially variable gene expression features instead of principal components of all genes, and (iii) SPRUCE allows for more flexible spatial correlation patterns compared to the global smoothing approach implemented by BayesSpace.

Finally, we applied SPRUCE to an un-annotated breast cancer sample sequenced with the 10X Visium platform. Using a set of the 16 top SVGs across the tissue sample, we discovered 5 unique cell clusters within the tissue sample. These clusters were marked by unique gene expression profiles which allowed us to characterize the biological function of each cluster using existing literature. We discovered an interesting interactions between a cluster of tumor resistant cells and a cluster of highly cancerous cells - an interplay which may have important implications for understanding the dynamics of the tumor microenvironment in the context of breast cancer.

This work may be extended in a number of promising ways. While we presented a general framework for accommodating a variety of spatial patterns using spatially correlated random effects, one might encode more specific biological hypotheses into the spatial component of the model through alternative prior distributions on the mixture component labels. Finally, while we developed SPRUCE for the quickly developing field of spatial transcriptomics, the model is generally applicable to multivariate data that feature spatial correlation across areal units.

## Supporting information

Web Appendix

## Acknowledgements

This work has been supported through grants from the National Institute of General Medical Sciences (R01 GM122078), National Cancer Institute grant (R21 CA209848), National Institute on Drug Abuse (U01 DA045300).

## Supplementary Materials

Web Appendix A, containing the proof of Proposition 1 discussed in Section 3, and Web Appendix B containing the MCMC algorithm discussed throughout Section 3 are available through *bioRxiv*.

## Notes

### Competing Interest Statement

The authors have declared no competing interest.

