## Supplementary material for "A Bayesian Multivariate Mixture Model for Spatial Transcriptomics Data": Web Appendix

#### Web Appendix A: Proof of Proposition 1

**Proposition 1** *Let $\pi_{ik}$ follow the multinomial logit model defined in equation (6) of the manuscript, and let $\psi_{ik}$ have a univariate intrinsic CAR prior as defined in equation (7) of the manuscript. Under Pólya–Gamma data augmentation, the full conditional distribution of $\psi_{ik}$ is $N(m_{ik}, V_{ik})$, where*

$$m_{ik} = \frac{\frac{1}{m_i}\sum_{l\in\delta_i}\psi_{lk} + U^*_{ik}}{\frac{m_i^2}{\nu_k^2} + \frac{1}{\omega_{ik}^2}}, \textit{ and } V_{ik} = \frac{1}{\frac{m_i^2}{\nu_k^2} + \frac{1}{\omega_{ik}^2}}, \tag{1}$$

*where $U^*_{ik} = \frac{U_{ik}-1/2}{\omega_{ik}} + c_{ik} - \mathbf{w}_i^T\boldsymbol{\rho}_k$, $U_{ik}$ is an indicator equal to 1 if $z_i = k$ and 0 otherwise, $c_{ik} = log(\sum_{h\neq k}^K exp(\mathbf{w}_i^T\boldsymbol{\rho}_h + \psi_{ih}))$, and $\omega_{ik} \sim PG(1, 0)$.*

*Proof.* *We first demonstrate the proof for the spot-level model. The full conditional distribution of $\psi_{ik}$, denoted $p(\psi_{ik}|\cdots)$, may be expressed as*

$$\begin{aligned} p(\psi_{ik}|\cdots) &= p(\psi_{ik}|\psi_{-ik}, \mathbf{z}, ...) = \frac{p(\psi_{ik}, \psi_{-ik}, \mathbf{z}, ...)}{p(\psi_{-ik}, \mathbf{z}, ...)} \\ &\propto \underbrace{p(\psi_{ik}|\psi_{-ik})}_{\textit{CAR prior}}\underbrace{p(\mathbf{z}|\psi_{ik}, \psi_{-ik}, ...)}_{\textit{likelihood}} \\ &\propto \left[N\left(\frac{1}{m_i}\sum_{l\in\delta_i}\psi_{lk}, \frac{\nu_k^2}{m_i}\right)\right]\prod_{i=1}^n\prod_{k=1}^K \pi_{ik}^{U_{ik}}, \end{aligned}$$

*where $U_{ik}$ is an indicator equal to 1 if $z_i = k$ and 0 otherwise. Given $\mathbf{U}_k = (U_{1k}, ..., U_{nk})^T$, we may re-parameterize the model for $\pi_{ik}$ as*

$$\begin{aligned} \pi_{ik} &= P(U_{ik} = 1) \\ &= \frac{exp(\mathbf{w}_i^T\boldsymbol{\rho}_k + \psi_{ik})}{\sum_{h=1}^K exp(\mathbf{w}_i^T\boldsymbol{\rho}_h + \psi_{ih})} \\ &= \frac{exp(\mathbf{w}_i^T\boldsymbol{\rho}_k + \psi_{ik})}{\sum_{h\neq k}^K exp(\mathbf{w}_i^T\boldsymbol{\rho}_h + \psi_{ih}) + exp(\mathbf{w}_i^T\boldsymbol{\rho}_k + \psi_{ik})}, \end{aligned}$$

*where dividing through by $\sum_{h\neq k}^K exp(\mathbf{w}_i^T\boldsymbol{\rho}_h + \psi_{ih})$ gives*

$$\pi_{ik} = \frac{exp(\mathbf{w}_i^T\boldsymbol{\rho}_k + \psi_{ik} - c_{ik})}{1 + exp(\mathbf{w}_i^T\boldsymbol{\rho}_k + \psi_{ik} - c_{ik})} = \frac{exp(\gamma_{ik})}{1 + exp(\gamma_{ik})},$$

*where $c_{ik} = log(\sum_{h\neq k}^{K} exp(\mathbf{w}_i^T \boldsymbol{\rho}_h + \psi_{ih}))$ and $\gamma_{ik} = \mathbf{w}_i^T \boldsymbol{\rho}_k + \psi_{ik} - c_{ik}$. Now, we notice that with respect to $\psi_{ik}$, the likelihood contribution may be simplified as follows.*

$$
\begin{aligned}
p(\psi_{ik}|\cdots) &\propto p(\psi_{ik}|\psi_{-ik}) \left(\frac{e^{\gamma_{ik}}}{1+e^{\gamma_{ik}}}\right)^{U_{ik}} \left(\frac{1}{1+e^{\gamma_{ik}}}\right)^{1-U_{ik}} \\
&= p(\psi_{ik}|\psi_{-ik}) \left[\frac{\left(\frac{e^{\gamma_{ik}}}{1+e^{\gamma_{ik}}}\right)^{U_{ik}} \left(\frac{1}{1+e^{\gamma_{ik}}}\right)}{\left(\frac{1}{1+e^{\gamma_{ik}}}\right)^{U_{ik}}}\right] \\
&= p(\psi_{ik}|\psi_{-ik}) \left[\left(\frac{\frac{e^{\gamma_{ik}}}{1+e^{\gamma_{ik}}}}{\frac{1}{1+e^{\gamma_{ik}}}}\right)^{U_{ik}} \left(\frac{1}{1+e^{\gamma_{ik}}}\right)\right] \\
&= p(\psi_{ik}|\psi_{-ik}) \frac{(e^{\gamma_{ik}})^{U_{ik}}}{1+e^{\gamma_{ik}}},
\end{aligned}
$$

*where the likelihood is now in the logistic form, which Polson et al. (2013) showed can be written as a scale mixture of normals with Pólya–Gamma precision terms $\omega_{ik} \sim PG(1,0)$. Thus, we have*

$$
\begin{aligned}
p(\psi_{ik}|\cdots) &\propto p(\psi_{ik}|\psi_{-ik}) \left[e^{(U_{ik}-1/2)\gamma_{ik}} \int_0^\infty e^{-\frac{\omega_{ik}\gamma_{ik}^2}{2}} p(\omega_{ik}) d\omega_{ik}\right] \\
&= p(\psi_{ik}|\psi_{-ik}) exp\{(U_{ik}-1/2)\gamma_{ik} - \omega_{ik}\gamma_{ik}^2/2\} \\
&\propto p(\psi_{ik}|\psi_{-ik}) exp\left\{-\frac{1}{2}\left(\frac{(U_{ik}^* - \psi_{ik})^2}{\omega_{ik}^2}\right)\right\},
\end{aligned}
$$

*where $U_{ik}^* = \frac{U_{ik}-1/2}{\omega_{ik}} + c_{ik} - \mathbf{w}_i^T \boldsymbol{\rho}_k$. Following standard results from Bayesian normal models (Hoff, 2009) we find the full conditional of $\psi_{ik}$ is $N(m_{ik}, V_{ik})$, as defined in equation (1).*

### Web Appendix B: MCMC Algorithm

1. *Update multivariate skew-normal outcome model parameters $\boldsymbol{\mu}_k$, $\boldsymbol{\xi}_k$, and $\boldsymbol{\Sigma}_k$.* For $k = 1, ..., K$:
   (a) Update $\boldsymbol{\mu}_k$:
      i. Define $n_k = \sum_{i=1}^n I_{z_i=k}$ as the number of spots in mixture component $k$.
      ii. Define $\mathcal{Z}_k$ as the set of all spot indices assigned to mixture component $k$.
      iii. Define $\mathbf{Y}_k$ as the $n_k \times g$ matrix of gene expression values for mixture component $k$. Similarly define $\mathbf{t}_k$ as the length $n_k$ vector of truncated normal random effects for mixture component $k$ and define $\boldsymbol{\Phi}_k$ as the $n_k \times g$ matrix with rows $\boldsymbol{\phi}_1^T, ..., \boldsymbol{\phi}_{n_k}^T$.
      iv. Define $\mathbf{E}_k = (\mathbf{Y}_k - \boldsymbol{\Phi}_k - \mathbf{t}_k^T \boldsymbol{\xi}_k)$ and $\bar{\mathbf{e}}_k$ as the column means of $\mathbf{E}_k$.
      v. Update $\boldsymbol{\mu}_k$ from its $\text{N}_g(\boldsymbol{\mu}_{nk}, \mathbf{V}_{nk})$ full conditional, where $\boldsymbol{\mu}_{nk} = \mathbf{V}_{nk}(\mathbf{V}_{0k}^{-1}\boldsymbol{\mu}_{0k} + n_k \boldsymbol{\Sigma}_k^{-1} \bar{\mathbf{e}}_k)$ and $\mathbf{V}_{nk} = (\mathbf{V}_{0k}^{-1} + n_k \boldsymbol{\Sigma}_k^{-1})^{-1}$.
   (b) Update $\boldsymbol{\xi}_k$:
      i. Define $\mathbf{M}_k$ as the $n_k \times g$ matrix with each row equal to $\boldsymbol{\mu}_k^T$.
      ii. Re-define $\mathbf{E}_k = \mathbf{t}_k \circ (\mathbf{Y}_k - \mathbf{M}_k - \boldsymbol{\Phi}_k)$, where "$\circ$" denotes the Hadamard product, and let $\bar{\mathbf{e}}_k$ be the $g \times 1$ vector of columns means of $\mathbf{E}_k$.
      iii. Update $\boldsymbol{\xi}_k$ from its $\text{N}_g(\boldsymbol{\xi}_{nk}, \mathbf{X}_{nk})$ full conditional, where $\boldsymbol{\xi}_{nk} = \mathbf{X}_{nk}(\mathbf{X}_{0k}^{-1}\boldsymbol{\xi}_{0k} + n_k \boldsymbol{\Sigma}_k^{-1} \bar{\mathbf{e}}_k)$ and $\mathbf{X}_{nk} = (\mathbf{X}_{0k}^{-1} + (\sum_{i \in \mathcal{Z}_k} t_i^2)\boldsymbol{\Sigma}_k^{-1})^{-1}$.
   (c) Update $\boldsymbol{\Sigma}_k$:
      i. Re-define $\mathbf{E}_k = (\mathbf{Y}_k - \mathbf{M}_k - \boldsymbol{\Phi}_k - \mathbf{t}_k^T \boldsymbol{\xi}_k)$.
      ii. Update $\boldsymbol{\Sigma}_k$ from its $\text{IW}(\nu_{nk}, \mathbf{S}_{nk})$ full conditional, where $\nu_{nk} = \nu_{0k} + n_k$ and $\mathbf{S}_{nk} = \mathbf{S}_{0k} + \mathbf{E}_k^T \mathbf{E}_k$.
2. *Update multivariate skew-normal conditional representation random effects $t_i$.* For $i = 1, ..., n$:

(a) Given $z_i = k$, define $A_k = (1 + \boldsymbol{\xi}_k^T \boldsymbol{\Sigma}_k^{-1} \boldsymbol{\xi}_k)^{-1}$ and define $a_{ik} = A_k(\boldsymbol{\xi}_k^T \boldsymbol{\Sigma}_k^{-1}(\mathbf{y}_i - \boldsymbol{\mu}_k - \boldsymbol{\phi}_i))$.

(b) Update $t_i$ from $\mathrm{N}_{[0,\infty)}(a_{ik}, \sqrt{A_k})$.

3. *Update outcome model multivariate CAR random effects* $\boldsymbol{\phi}_i$. For $i = 1, ..., n$:

   (a) Given $z_i = k$, update $\boldsymbol{\phi}_i$ from its $g-$dimensional normal full conditional, where $\mathrm{E}(\boldsymbol{\phi}_i|...) = (\boldsymbol{\Sigma}_k^{-1} + m_i \boldsymbol{\Lambda})^{-1}(\boldsymbol{\Sigma}_k^{-1}(\mathbf{y}_i - \boldsymbol{\mu}_k - t_i \boldsymbol{\xi}_k) + \boldsymbol{\Lambda}^{-1} \sum_{l \in \delta_i} \boldsymbol{\phi}_l)$ and $\mathrm{Cov}(\boldsymbol{\phi}_i|...) = (\boldsymbol{\Sigma}_k^{-1} + m_i \boldsymbol{\Lambda})^{-1}$.

4. *Update multinomial regression CAR random effects* (See Proposition 1). For $i = 1, ..., n$ and $k = 1, ..., K$, the full conditional distribution of $\psi_{ik}$ is $\mathrm{N}(m_{ik}, V_{ik})$, where

$$m_{ik} = \frac{\frac{1}{m_i} \sum_{l \in \delta_i} \psi_{lk} + U_{ik}^*}{\frac{m_i^2}{\nu_k^2} + \frac{1}{\omega_{ik}^2}}, \text{ and } V_{ik} = \frac{1}{\frac{m_i^2}{\nu_k^2} + \frac{1}{\omega_{ik}^2}}, \tag{2}$$

   where $U_{ik}^* = \frac{U_{ik} - 1/2}{\omega_{ik}} + c_{ik} - \mathbf{w}_i^T \boldsymbol{\rho}_k$, $U_{ik}$ is an indicator equal to 1 if $z_i = k$ and 0 otherwise, $c_{ik} = \log(\sum_{h \neq k}^K \exp(\mathbf{w}_i^T \boldsymbol{\rho}_h + \psi_{ih}))$, and $\omega_{ik} \sim \mathrm{PG}(1, 0)$.

5. *Update multinomial regression mixing weight parameters* $\boldsymbol{\rho}_k$ *and latent variables* $\omega_{ik}$. For $k = 1, ..., K - 1$:

   (a) For $i = 1, ..., n$, update PG latent variables $\omega_{ik}$ from $\mathrm{PG}(1, \eta_{ik})$, where $\eta_{ik} = \mathbf{w}_i^T \boldsymbol{\rho}_k - c_{ik}$.

   (b) Compute $\mathbf{R}_{nk} = (\mathbf{R}_{0k}^{-1} + \mathbf{W}^T \mathbf{O}_k \mathbf{W})$, where $\mathbf{O}_k$ is the diagonal matrix with entries $(\omega_{1k}, ..., \omega_{nk})$, $\mathbf{W}$ is the $n \times p$ matrix of covariates with rows $\mathbf{w}_1^T, ..., \mathbf{w}_n^T$.

   (c) Compute $\boldsymbol{\rho}_{nk} = \mathbf{R}_{nk}(\mathbf{R}_{0k}^{-1} \boldsymbol{\rho}_{0k} + \mathbf{W}^T \mathbf{O}_k \mathbf{U}_k^*)$, where $\mathbf{U}_k^* = \left(\frac{U_{1k} - 1/2}{\omega_{1k}} + c_{1k}, ..., \frac{U_{nk} - 1/2}{\omega_{nk}} + c_{nk}\right)$.

   (d) Update $\boldsymbol{\rho}_k$ from $\mathrm{N}_p(\boldsymbol{\rho}_{nk}, \mathbf{R}_{nk})$.

6. *Update mixture component labels* $z_1, ..., z_n$. For $i = 1, ..., n$:

   (a) Compute the probability of spot $i$ belonging to cluster $k$ under current values of model parameters. For $k = 1, ..., K$, compute $P_{ik} = \mathrm{dnorm}(\mathbf{y}_i; \boldsymbol{\mu}_k + \boldsymbol{\phi}_k + t_i \boldsymbol{\xi}_k, \boldsymbol{\Sigma}_k)$.

   (b) Compute $\pi_{ik} = \frac{\exp(\mathbf{w}_i^T \boldsymbol{\rho}_k + \psi_{ik})}{\sum_{h=1}^K \exp(\mathbf{w}_i^T \boldsymbol{\rho}_h + \psi_{ih})}$.

   (c) Compute $\mathrm{P}(z_i = k|...) = \frac{P_{ik} \pi_{ik}}{\sum_{h=1}^K P_{ih} \pi_{ih}}$.

   (d) Update $z_i$ from Categorical$\{\mathrm{P}(z_i = 1|...), ..., \mathrm{P}(z_i = 1|...)\}$.

7. *Re-map mixture component labels to protect against label switching.*

   (a) Define $\mathrm{ord}(\mathbf{z})$ as the function to return a length $K$ vector containing the order in which each unique mixture component label appears in $\mathbf{z}$. E.g., in `R` this is the `unique()` function.

   (b) Initialize $\mathbf{z}^*$ as an empty length $n$ vector for the re-mapped mixture component labels.

   (c) For $k = 1, ..., K$, let $\mathbf{z}^*[\mathbf{z} = \mathrm{ord}(\mathbf{z})[k]] = k$.
